# MERIT: a Mutation Error Rate Identification Toolkit for Ultra-deep Sequencing Applications

**DOI:** 10.1101/184291

**Authors:** Mohammad Hadigol, Hossein Khiabanian

## Abstract

Rapid progress in high-throughput sequencing (HTS) has enabled the molecular characterization of mutational landscapes in heterogeneous populations and has improved our understanding of clonal evolution processes. Analyzing the sensitivity of detecting genomic mutations in HTS requires comprehensive profiling of sequencing artifacts. To this end, we introduce MERIT, designed for in-depth quantification of erroneous substitutions and small insertions and deletions, specifically for ultra-deep applications. MERIT incorporates an all-inclusive variant caller and considers genomic context, including the nucleotides immediately at 5′ and 3′, thereby establishing error rates for 96 possible substitutions as well as four singlebase and 16 double-base indels. We apply MERIT to ultra-deep sequencing data (1,300,000×) and show a significant relationship between error rates and genomic contexts. We devise an *in silico* approach to determine the optimal sequencing depth, where errors occur at rates similar to those of true mutations. Finally, we assess nucleotide-incorporation fidelity of four high-fidelity DNA polymerases in clinically relevant loci, and demonstrate how fixed detection thresholds may result in substantial false positive as well as false negative calls.

## Introduction

Advances in high-throughput sequencing (HTS) technologies have revolutionized the genomic, transcriptomic, and epigenomic characterization of biological states. HTS platforms produce large amounts of sequencing reads at relatively low cost. However, their high rates of sequencing artifacts have hindered profiling heterogenous populations such as those in tumor samples, where in addition to genomically diverse cancer cells, contaminating normal cells may also be present. Comprehensive error profiling and mutation detection sensitivity analysis has direct implications for predicting evolutionary trajectories that may arise from such heterogeneity. For example, since cancer’s eventual evolution to a therapy-resistant disease is often associated with the natural selection of pre-existing resistant clones, detecting somatic genomic variations that exist in small cell populations prior to treatment is pertinent for prevention of therapy-resistant relapsed disease. [1, 2]

HTS errors are dominated by misreading a base within the instrument or nucleotide misincorporations during library enrichment with polymerase chain reaction (PCR). Specifically, DNA damaging factors such as spontaneous deamination, presence of oxidized bases in cells in addition to *ex vivo* oxidation during DNA extraction [3], or short-lived high temperatures during acoustic shearing [4] have been associated with high polymerase error rates [5]. PCR-free library preparation [6, 7], sophisticated DNA barcoding [4, 8, 9], and overlapping-read design [10, 11, 12] have been proposed to reduce rates of error. Despite varying success of these methods, complete elimination of HTS errors is not possible; therefore, their exhaustive analysis is critical for estimating and improving sensitivity in calling somatic variants.

Here, we introduce MERIT (Mutation Error Rate Identification Toolkit), a comprehensive pipeline designed for in-depth quantification of erroneous HTS calls, specifically for ultra-deep applications. MERIT considers the genomic context of errors and shows a significant relationship between error rates and their sequence contexts. In addition to observing more than three orders of magnitude difference between transition and transversion error rates, we identify variations of more than 130-fold within each error type at high sequencing depths. We also propose an *in silico* depth reduction approach to provide insights on estimating optimal depth - where sequencing errors exist at rates similar to those of true mutations. Finally, we describe an assay for detailed assessment of nucleotide-incorporation fidelity for four high-fidelity DNA polymerase molecules.

## Results

### MERIT: a comprehensive error rate estimator

Differential rate of substitution errors in HTS has been attributed to common DNA damaging events such as deamination and oxidation during library preparation. Such events often lead to higher rates of transitions versus transversions [13, 14, 15, 16, 17] or increased number of errors in specific genomic contexts.These differences can be more pronounced at higher sequencing depths and directly impact the sensitivity for detecting true mutations. MERIT, primarily designed for profiling ultra-deep data (depths >5,000×), provides a comprehensive toolkit to identify, characterize, and quantify various sources of HTS error.

Comparative performance analysis of the commonly used HTS variant callers [18, 19, 20] suggests a significant disagreement between their identified variants [21, 22, 23]. These differences are mainly rooted in each pipeline’s specific filtering and statistical methodology. Moreover, since the goal of these algorithms is to distinguish true mutations from artifacts, they do not report all the variants required for a precise quantification of sequencing noise. To overcome this limitation, MERIT uses SAMtools to identify all positions with alternate alleles from the aligned, indexed sequencing reads. It then probes SAMtools mpileup data to extract the reference and alternate alleles’ depths, Phred quality, and position-in-read information for all variants, even when they are present in only a single read amongst tens of thousands. Finally, it obtains the genomic context of the variants from the reference genome, including the nucleotides immediately at their 5′ and 3′, and estimates error rates for 96 possible single nucleotide substitutions as well as four single-base and 16 double-base insertions/deletions (indels). An optional annotation step is also available. Details of MERIT’s workflow are shown in Figure 1 and described in Methods.

**Figure 1:**
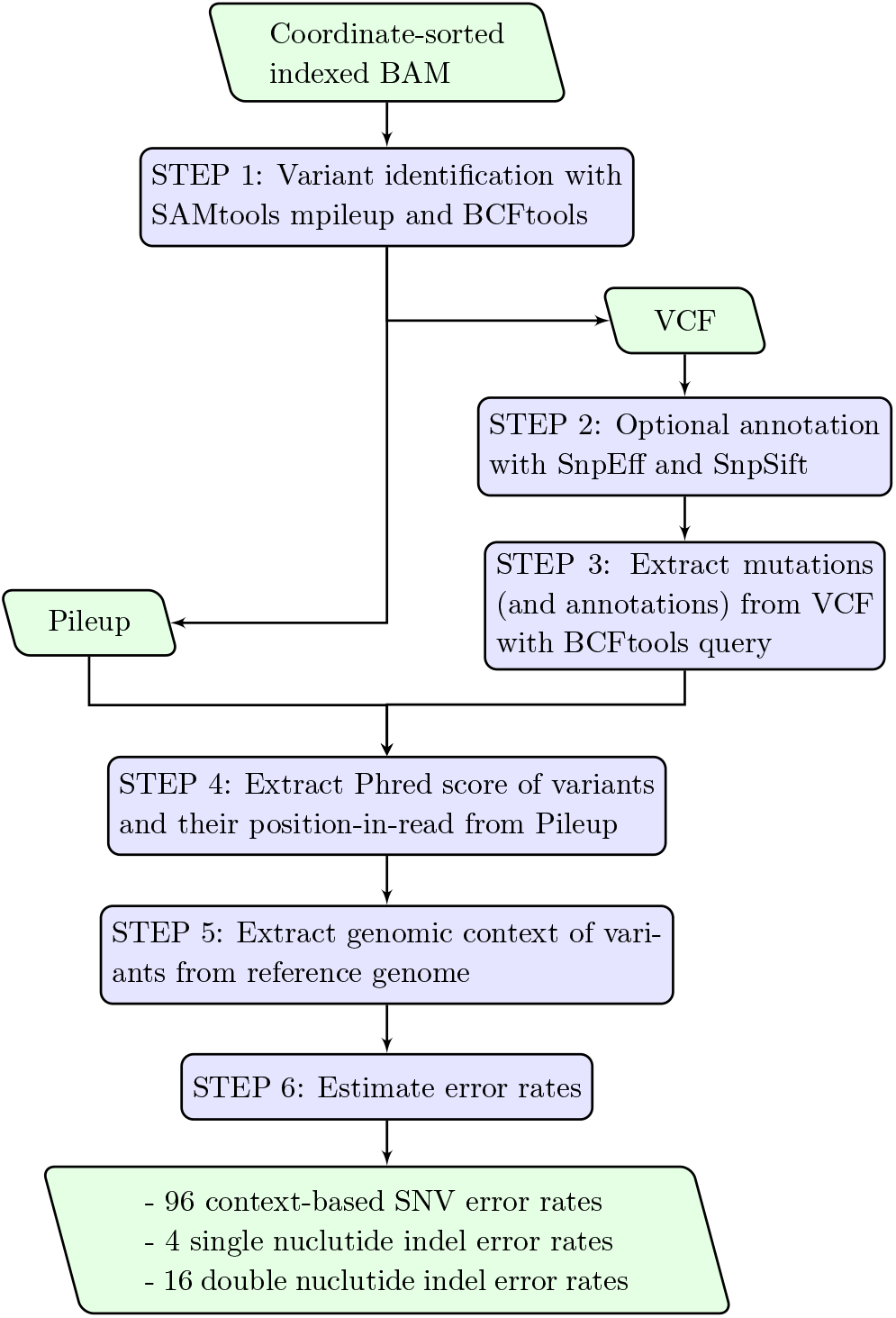
MERIT’s workflow, developed for comprehensive characterization of the sequencing error rate in ultra-deep HTS applications.

### Ultra-deep sequencing data set

We generated ultra-deep sequencing data for a HapMap sample (NA19240) targeting four commonly mutated loci in the *TP53* and *SF3B1* genes. We used four high-fidelity DNA polymerase enzymes - NEBNext^®^ High-Fidelity 2X PCR Master Mix (Hi-Fi 2X), NEBNext^®^ Ultra^™^ II Q5^®^ Master Mix (Ultra II), KAPA HiFi PCR kits with ReadyMix (KAPA), and Invitrogen^™^ Platinum^™^ SuperFi^™^ DNA polymerase (SuperFi) - for PCR amplification. Custom primers for amplicon-based paired-end reads were designed such that a large portion of the two reads for each DNA fragment (R1 and R2) overlapped (Tables S1 and S3, SI Materials). Because we used a single, and therefore homogenous, sample, it followed that all detected variants were errors accumulated in library preparation or during sequencing.

### Impact of merging reads on context-specific error correction

Independent analysis of R1 and R2 reads at 1,300,000 × indicated significant difference in estimated error rates across 96 possible sequence contexts (Fig. 2). High error rates and low Phred quality scores observed in R2 relative to R1 may be associated with sequencing errors caused by misreading a base, attributed to image analysis biases [5] or phasing/pre-phasing [17]. These sequencing errors that dominated the R1 and R2 profiles can be distinguished from polymerase errors by merging the overlapped paired-end reads [24, 11, 25]. Merging, however, cannot eliminate errors randomly accumulated during the amplification processes and present in both reads. In contrast to higher rates of transversion versus transition errors in paired-end reads (Fig. 3A), the remaining PCR-related errors in the merged reads are dominated by transitions, often with high Phred quality scores (Fig. 3B). MERIT provides further insight for profiling these errors, which are the main hurdle in distinguishing real mutations from sequencing noise:

- Merging R1 and R2 reads lowered all the context-specific error rates. The highest reduction in rate was observed for GTA>GGA transversions (5,025±2,794×) while GCG>GTG transition errors only improved by a factor of 1.22±0.07×. Moreover, these improvements were context-specific. For example, T>A transversion in GTA trinucleotides showed substantial reduction (568±249×) compared to those in CTAs (1.43±0.31×).
- Transition errors occurred at higher rates relative to transversions, in agreement with previous reports [13, 14, 15, 16, 17]. This difference was pronounced further when errors were classified based on their context, denoting a rate of 1.29±0.04×10^−3^ [error/base] for GCG>GTG versus that of 2.17±0.92×10^−6^ [error/base] for GTA>GAA (Fig. 3B). MERIT also revealed considerable variation within each substitution type. For example, T>G transversions in GTCs occurred 133.5±65.9× more often than those in ATAs. Similarly, C>T transitions in GCGs were observed at 73.8±10.5× higher rate than those in TCTs (Fig. 3B).
- The rate of C>A errors in ACCs was the highest of all such transversions. These errors are linked to the conversion of guanine to 8-oxoG resulting in mismatched pairing with adenine [26, 27]. Oxidation of guanine to 8-oxoG happens naturally in living cells and can be increased by DNA damaging factors such as acoustic shearing [28].
- Merging R1 and R2 can correct for the low quality erroneous bases associated with sequencing errors. Our analysis suggests that such sequencing errors can be identified and eliminated based on their quality, when merging the reads is not possible (e.g., in hybrid-capture-based sequencing where read pairs are not designed to necessarily overlap).

**Figure 2:**
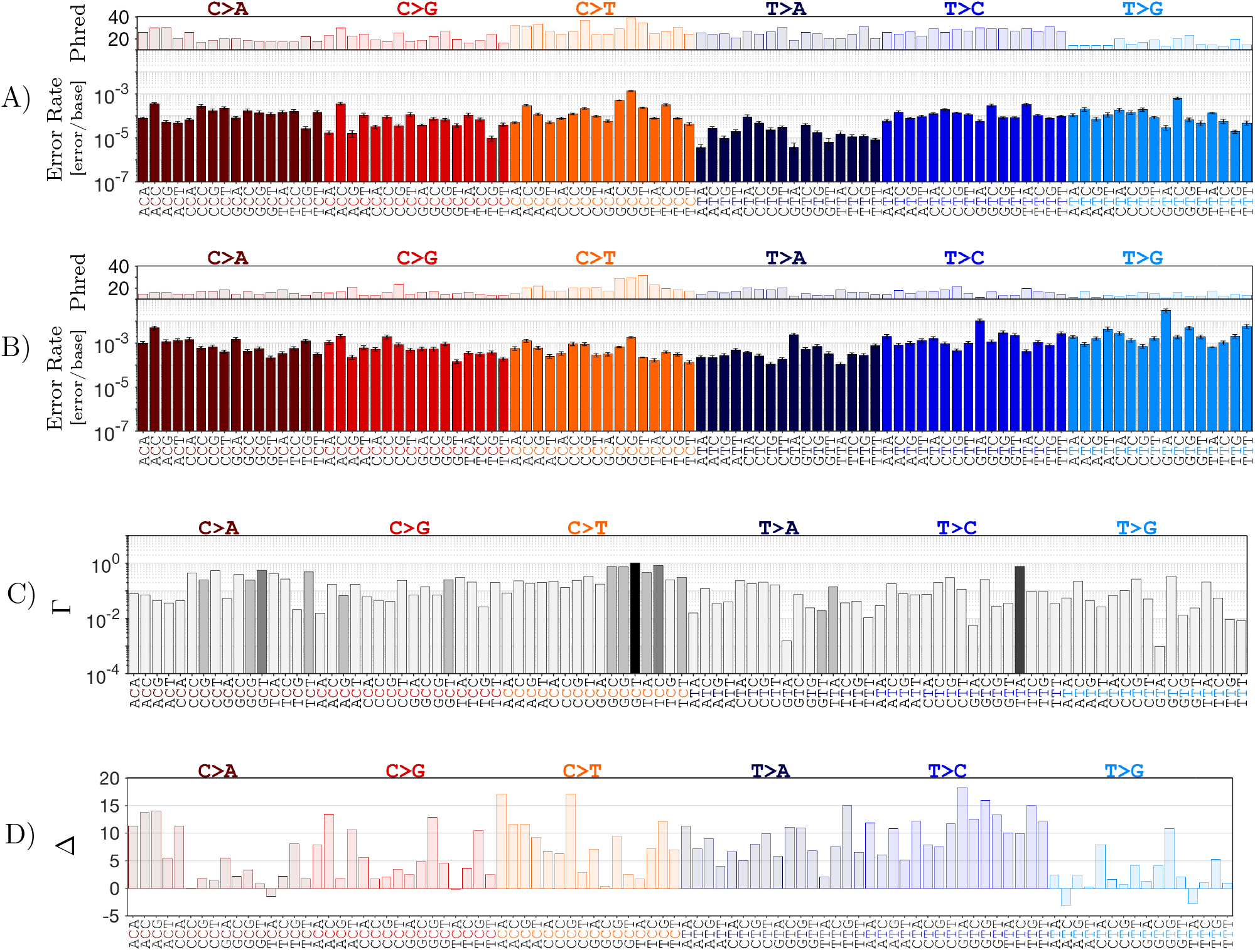
Estimated context-specific substitution error rates for polymerase Hi-Fi 2X. A) R1 reads. B) R2 read. C) Γ, the ratio of error rate in R1 over R2. P-values are computed by performing a two-tailed z-test: p-value > 0.05 ◼, 10^−5^ < p-value < 0.05 ■, 10^−10^ < p-value < 10^−5^ ■, 10^−50^ < p-value < 10^−10^ ■, p-value < 10^−50^ ■. D) Δ, the difference between their corresponding Phred quality scores. We reduced the depth of paired-end reads to approximately 1,300,000× through an *in silico* depth reduction experiment. Results are obtained by averaging over 100 independent samples to establish error bars which indicate one standard deviation from the average.

**Figure 3:**
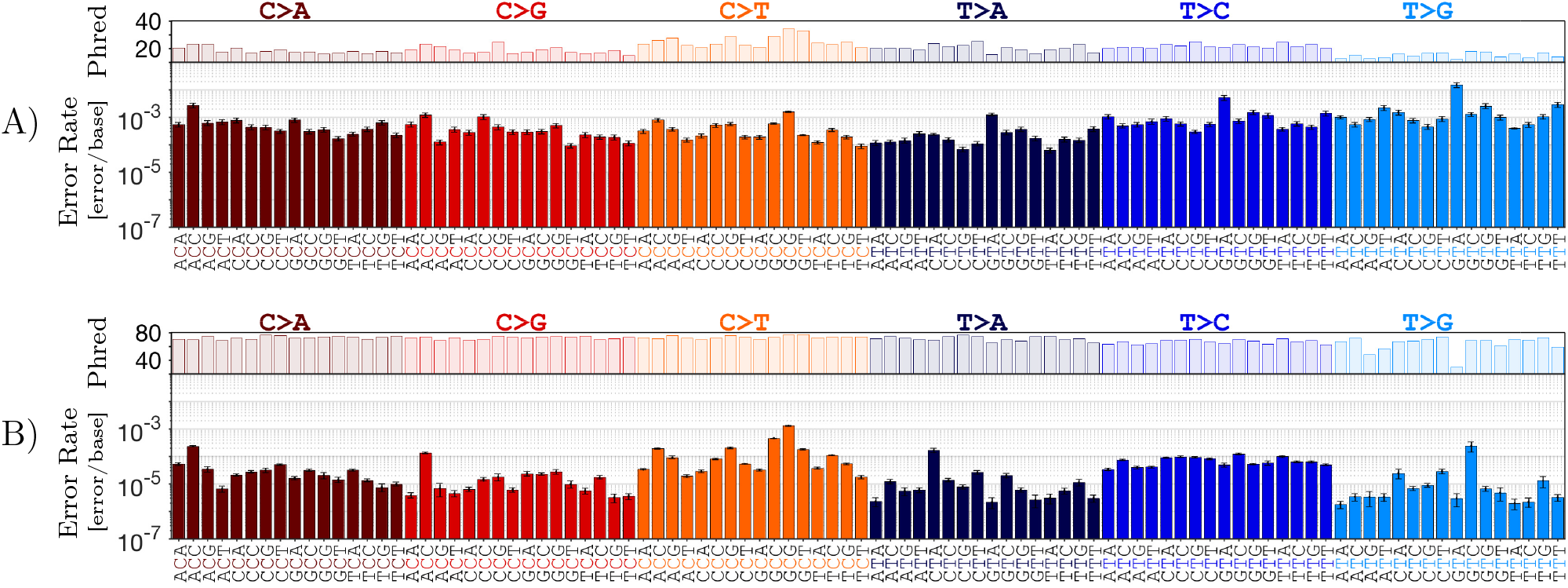
Merging of overlapped PE reads reduces context-specific error rates. A) R1 and R2 reads. B) merged reads. Depth of merged reads for polymerase Hi-Fi 2X were reduced in *silico* to approximately 650,000×. Error bars indicate one standard deviation from the average of 100 independent sub-samples.

### Effect of mutation context on amino acid variations

In a single codon, the context-specific rate of error for each base change directly affects the sensitivity of detecting the resulting amino acid variation. Our data indicated that the most commonly mutated residues in *TP53* and *SF3B1* were often more prone to errors and hence comparatively less likely to be distinguished from sequencing errors. For example, in *TP53*, R248Q and R248W are among the most common mutations found in cancer patients [29]. The transition base changes that result in these mutations could be confounded by the HTS errors at an 8-fold higher rate than the transversion alterations that lead to R248L, and 55-fold higher than those that lead to R248G (Fig. 4A). Similarly, the K700E mutation in *SF3B1* is the most frequently mutated residue in the gene’s exon 10 [30, 31]; it results from a T>C mutation in a TTC trinucleotide that showed the highest rate of error for a non-synonymous amino acid change in its codon (4.74±0.42×10^−5^ [error/base]). In contrast, the comparatively rarer I704F mutation — a T>A in a ATG — had one of the lowest rates of error in its respective codon (5.15±1.13×10^−6^ [error/base]; Fig. 4B). K700E’s 9-fold higher rate of error than that of I704F indicated marked reduction in its relative detection sensitivity.

**Figure 4:**
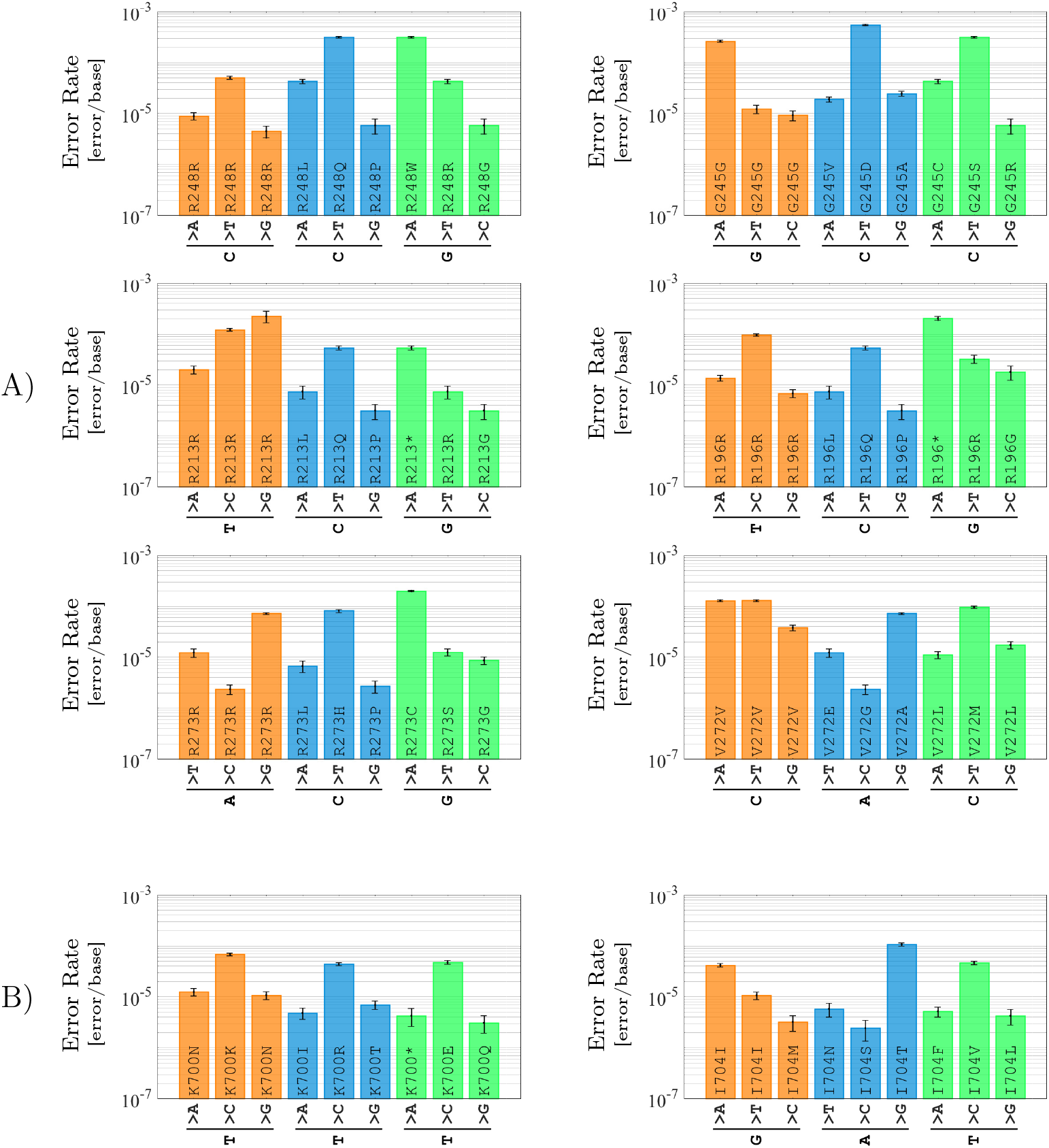
Significant variation in error rates for possible amino acid changes at individual codons. A) Six frequently mutated residues in the *TP53* gene. B) Two hotspot residues in the *SF3B1* gene. The higher the rate of error for a specific base change, the lower the power to distinguish true mutations from sequencing artifacts at its position. Here, the error rates represent the amplification by the Hi-Fi 2X polymerase. Error bars represent one standard deviation from the mean of 100 independent sub-samples.

### Optimal sequencing depth

Insufficient sequencing depth reduces the sensitivity of detecting variants and leads to loss of statistical significance for a confident variant calling [32]. Consequently, sequencing at higher depths is expected to provide robust error rate estimates and improved sensitivities in detecting true mutations. Accurate estimation of optimal sequencing depth, beyond which the inferred background error is not further reduced, not only provides a precise view of intrinsic limitations in HTS assays, but also leads to preserving time and resources by avoiding unproductive ultra-deep sequencing experiments.

To provide insight on optimal sequencing depth, we performed *in silico* experiments and estimated context-specific error rates as a function of depth. We randomly selected merged reads and constructed simulated sequencing data at depths ranging from 1,000 × to 700,000 ×, with 500 independent replicates at each depth to establish confidence intervals (Fig. 5). MERIT showed that the type of substitution error was an important determinant in estimating the optimal depth (Fig. 5A). The error rate estimates for all transitions as well as C>A transversions did not significantly change as sequencing depth increased beyond 200,000 ×; however, the inferred rates for the remaining transversions marginally improved at higher depths. More importantly, this analysis highlighted the importance of context-specific error profiling in determining detection sensitivity thresholds for true mutations. For example, at 5,000×, the corresponding error rates for all T>A errors, T>A errors in CTAs, and T>A errors in GTTs were 2.19±0.37×10^−4^ [error/base], 4.27±2.28×10^−4^ [error/base], and 1.96±0.02×10^−4^ [error/base], while at 700,000×, these rates were reduced to 2.02±0.73×10^−5^ [error/base], 2.5±0.99×10^−4^ [error/base], and 2.01±0.89×10^−6^ [error/base], respectively. Selecting a frequency threshold for these variants at 5,000× based on the general T>A rate may not yield significant number of false calls independent of their sequence contexts; however, at depths >5,000×, setting a threshold based on all T>A errors would lead to substantial false positive CTA>CAA and false negative GTT>GAT calls, as their corresponding error rates diverge at high depths, reaching a difference of two orders of magnitude at 700,000×.

**Figure 5:**
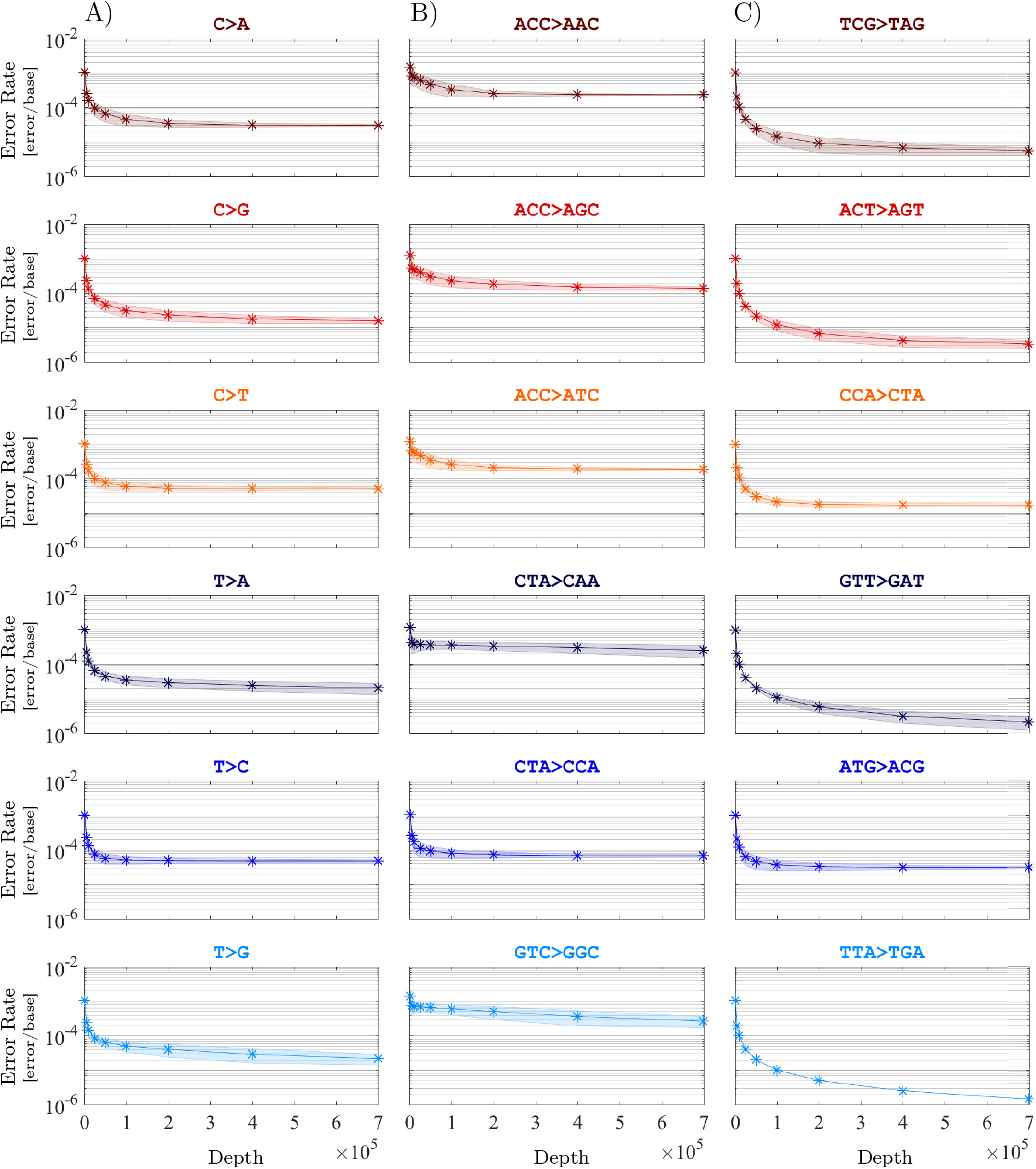
Substitution error rates are classified based on their type (column A) and context (columns B and C) at nine different depths: 1,000×, 5,000×, 10,000×, 25,0000×, 50,000×, 100,000×, 200,000×, 400,000×, and 700,000×. In *silico* depth reduction experiment was performed on merged reads, amplified by polymerase Ultra II to an average depth of 1,930,473×. The shaded areas are uncertainty bounds of one standard deviation around the average, derived from 500 independent sub-samples.

### DNA polymerase fidelity estimation

High-fidelity DNA polymerases - equipped with proofreading - result in fewer base misincorporations in PCR enrichment step, and thus, can reduce HTS error rates. The Hi-Fi 2X, Ultra II, KAPA, and SuperFi enzymes are marketed as high-fidelity polymerases, specifically designed for efficient amplification of complex templates such as those with GC-rich regions. Their providers have reported a fidelity 100 × better than wild-type Taq DNA polymerase [33, 34, 35, 36]. Several techniques such as forward mutation assay [37], denaturing gradient gel electrophoresis [38], and most recently, HTS-based techniques [39, 40, 41, 42] have been used to determine replication fidelity of DNA polymerases. Because different assays, quantification methods, and often descriptive units [42, 41] are used to report polymerase fidelity, comparing these methodologies is a challenging task and beyond the scope of this work. Instead, here, we used MERIT to emphasize on the importance of context-specific error estimation and provide a robust comparison of these commonly used enzymes.

We applied MERIT to the merged reads at equal depths of 650,000 ×, ensuring that the estimated fidelities were not affected by sequencing depth. When all errors were included in the analysis, global error rates suggested that these polymerases performed fairly similarly to each other, with the highest and lowest error rates belonging to KAPA and SuperFi enzymes, respectively. Specifically, the overall substitution error rates for Hi-Fi 2X, Ultra II, KAPA, and SuperFi were estimated at 2.66±0.21×10^−6^, 1.91±0.19×10^−6^, 6.95±0.54×10^−6^, and 1.76±0.25×10^−6^ [error/base/doubling], respectively (Fig. S1A).

Despite highly correlated error profiles of these polymerases (Fig. 6D), MERIT illustrated enzyme-specific biases beyond global error rates. For example, the overall substitution fidelity of SuperFi was found 3.95±0.65× better than that of KAPA’s; however, specific substitution fidelity differed widely. C>G errors of SuperFi were 6.88±2.16× less frequent than those of KAPA. In contrast, for C>A substitutions, SuperFi’s advantage over KAPA was reduced to only 1.85±0.30× (Fig. S1B). More unexpected differences could also be observed across 96 contexts (Fig. 6A). For example, TTA>TGA error rate of SuperFi is 132±35× lower than KAPA, while for GCG>GAG errors, KAPA performs just slightly better than SuperFi. Finally, using the data from multiple regions of the *TP53* and *SF3B1* genes, we found limited change in error profiles as the similarities between the genomic content of the amplified amplicons decreased as measured by their corresponding Kullback-Leibler distance (Fig. 7).

**Figure 6:**
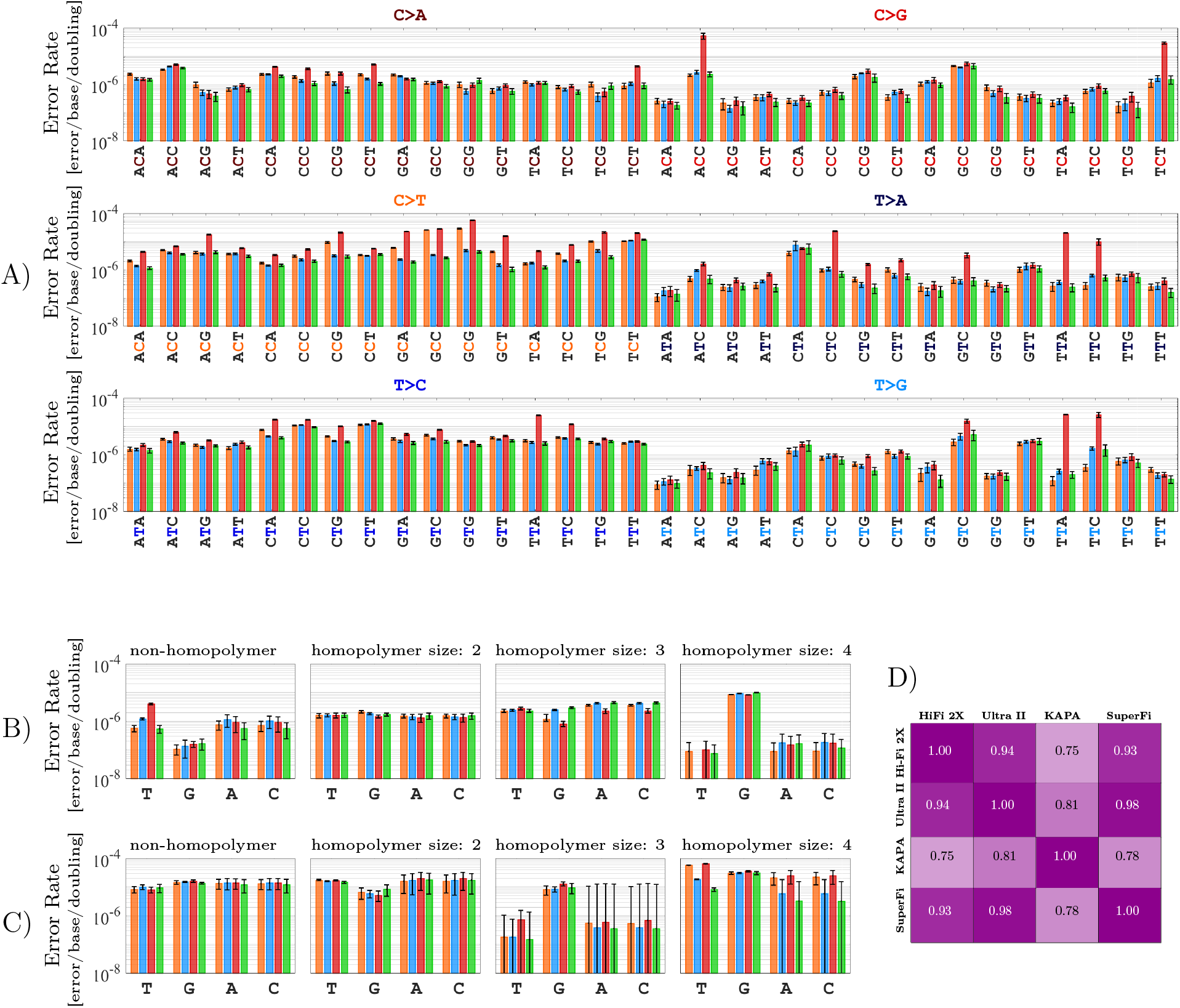
Estimating the fidelity of four polymerases, i.e., Hi-Fi 2X 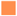, Ultra II 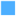, KAPA 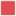, and SuperFi 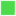. A) Context-specific substitutions. B) Single-base insertions. C) Single-base deletions. D) Spearman’s rank correlation coefficient between context-specific error profiles. Results are obtained by averaging over 100 independent samples to establish error bars, which indicate one standard deviation from the average.

**Figure 7:**
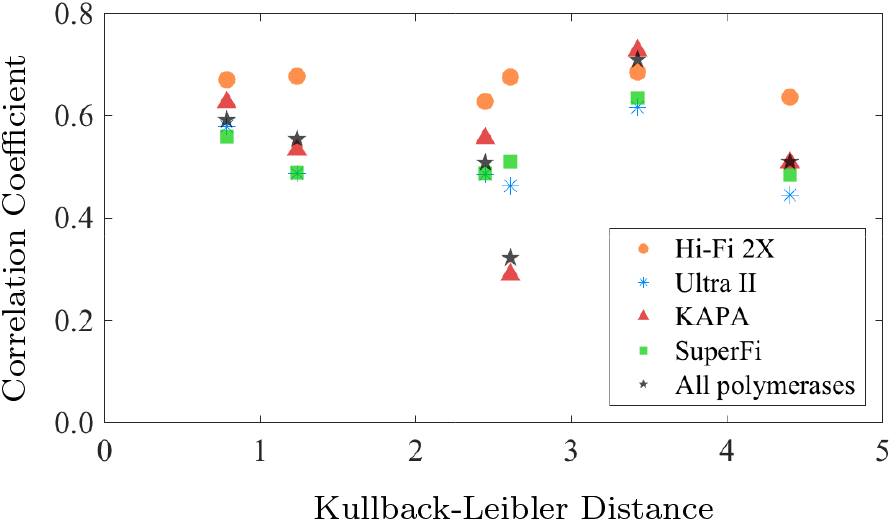
Spearman’s rank correlation coefficient between the context-specific error profiles of the targeted genes as a function of the symmetric Kullback-Leibler distance between their content profiles presented in SI Materials, Fig. S2.

## Discussion

Rapid progress in HTS technologies in addition to the development of novel library preparation methods have succeeded in reducing background sequencing noise, which has led to improving the sensitivity of detecting true mutations in heterogenous samples: PCR-free methods [43, 44] have forgone the bias associated with the polymerase base incorporation [6, 7]; DNA barcoding and overlapping read design approaches have minimized PCR-related errors [4, 8, 45, 46, 9]. However, these methods are bounded by their technical and theoretical limitations and often require large amounts of input DNA. Moreover, since sequencing error cannot be completely eliminated, comprehensive quantification of their profiles can highlight both the efficiency and the limitations of any methodology. In this manuscript, we introduce MERIT, which provides a platform-independent toolkit and illustrates the importance of estimating error rates in their genomic contexts, especially in ultra-deep HTS data.

Our results as applied to the Illumina platform confirm previous data on the differential rates of errors in paired-end sequencing reads [17], and indicate that merging the overlapping read pairs can notably correct errors that accumulate in sequencing instruments. We also provide a systematic approach for estimating the optimal sequencing depth for discovery of mutations that exist at very low abundances. Our data show that increasing sequencing depth may improve sensitivity for detecting some mutations based on their genomic context.

We also report the application of MERIT to ultra-deep sequencing data obtained from the amplification of multiple clinically relevant loci using four high-fidelity polymerase enzymes. Although there is limited variation in both the rates of error and dependence on the genomic content of the amplified region, our results indicate that profiling each polymerase’s misincorporations according to their genomic contexts has important clinical consequences. Specifically, we show that assigning a single allele frequency threshold to call mutations may result in substantial false positive as well as false negative variants. Not only are neighboring mutational hotspots in one gene affected with markedly different error rates, there is significant variation in the sensitivity of detecting common amino acid changes within each residue. These results suggest that some of these mutations may in fact be more prevalent at sub-clonal levels in disease populations than previously reported. Small mutated clones in the *TP53* and *SF3B1* genes, present in >0.1% of alleles, are shown to be strong predictors of poor survival and possible resistance to therapy in various neoplasms [47, 48, 49, 50]; thus, their detection at very low abundances is pertinent for patient care. Put together, our results strongly advocate mutation-specific variant calling approaches that go beyond estimating fixed detection thresholds for all variants [51].

## Methods

### DNA sample

HapMap NA19240 human genomic DNA (5 *µ*g) was obtained from Coriell, purified from immortalized lymphocytes using the Qiagen Autopure LS instrument in TE buffer (10 mM Tris, pH 8.0/1 mM EDTA) with concentration of 301 ng/L. Sample quality and concentration were assessed via Nanodrop and Qubit dsDNA assay before library preparation.

### HTS platform and assay optimization

Custom amplicon-based sequencing and library preparation were performed at GeneWiz (South Plainfield, NJ) using Illumina HiSeq2500 Rapid Run. Primers were designed using Primer3 [52] and were used along with Illumina partial adapters to target four loci in the *TP53* and *SF3B1* genes (SI Materials, Table S1) such that the paired-end reads significantly overlapped. The annealing temperature of 66°C was determined based on the product specificity and yield for all polymerases after performing gradient PCR optimization at eight different temperatures (SI Materials, Fig. S3).

### PCR amplification and indexing

Twenty PCR cycles were performed using the Hi-Fi 2X, KAPA, and SuperFi polymerases, and 16 cycles were performed using the Ultra II polymerase in the first round of amplification. The cycle numbers were determined after initial PCR amplification tests (Table S2, SI Materials) in order to obtain similar amount of DNA for each enzyme.The second round of PCR for multiplexing included seven cycles for all four polymerases. After each PCR amplification, AMPure Bead cleanup was performed. First, 0.4× ratio (20 *µ*L AMPure bead to 50 *µ*L PCR product) was used to remove gDNA and larger fragments (i.e., >600 bps). For the saved supernatant, additional 80 *µ*L AMPure Bead was added to bring the total to a 2× ratio. The beads were eluted with 22 *µ*L EB (10 mM Tris, pH 8.0). Qubit quantification and Bioanalyzer analysis were performed for quality assessment.

### Alignment and merging

Paired-end (PE) reads were cleaned of adapters and primers, and were aligned to the reference human genome hg19 assembly using the Burrows-Wheeler Aligner (BWA) tool [53]. PE reads that properly mapped to the targeted loci were then merged. In our merging scheme, if R1 and R2 reads did not match at a base, an N was assigned for that position. Read pairs with smaller than 50 base overlaps or with more than five mismatches were discarded. Phred quality score (*Q*) of a successfully merged locus was calculated as sum of the qualities in R1 and R2 reads since these are independent events; *Q* is given by *Q* = −10 log_10_ *p* where *p* is the probability that the base is called wrong. Merged reads were then mapped to the reference human genome, and were filtered so that they were uniquely mapped (BWA tags X0:1 and X1:0). Finally, in order to make a fair comparison between the error rate of merged and PE reads, only PE reads that were merged successfully and uniquely mapped were considered. Table S3 in SI Materials represents the average depth of merged and PE reads in different loci.

### In silico depth reduction

The sequencing assay was designed to obtain an average depth of >1,000,000× bp, but for some amplicons the average depth was substantially larger (Table S3, SI Materials). Therefore, an in *silico* depth reduction procedure was performed to reduce the high depths and more importantly, generate enough independent samples to estimate low error rates confidently. It should be noted that one of the main hurdles in error rate estimation of high fidelity polymerases via HTS is the lack of signal as errors occur infrequently with increased fidelity, hence, a large number of samples is required to accurately estimate errors. As performing ultradeep sequencing on a large number of samples is not cost-effective, alternatively, *in silico* samples at lower depths can be generated from one ultra-deep sequencing run by randomly selecting reads from the original raw sequencing data.

### Error rate estimation

Context-specific erroneous base calls at each locus were assumed to follow a binomial distribution. More precisely, the probability of a single nucleotide X_i_ with the genomic context *ZX_i_Z’, Z,Z’ ∈ {A,C,T,G}*, in a specific locus *i* being misread as *Y_i_*, i.e., *P_ZX_i_Z’→ZY_i_Z’_* followed

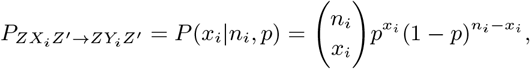

where *p* is the combined PCR and sequencing error rate and *n_i_* and *x_i_* are the total read depth and the number of erroneous calls at position *i*, respectively. Assuming a position-independent *p*, the probability of observing *m* instances of *ZXZ’−ZYZ’* error within each sample was then given by

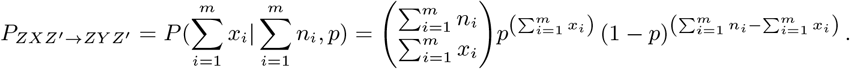

See Remark 1 in SI Materials on the sum of binomial random variables.

For the case of indels, a binomial model was used to describe the error rate as well, but instead of categorizing them based on their context, indels were classified based on the type of inserted/deleted base, as no differential error rates was observed for context-specific indels.

This estimated error rate has a unit of [error/base]. It is common to report the fidelity of polymerase enzymes as [error/base/doubling] in the literature where template doubling *d* is given by

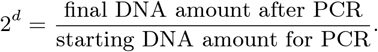

Since precise amounts of input and output DNA was known for our experiment in its second round of PCR, template doubling could be calculated and polymerase replication efficiency would be estimated as the ratio of template doubling d over the number of PCR cycles performed (Table S4). Next, to obtain the total amount of template doubling after performing two rounds of PCR amplification, the total number of PCR cycles were multiplied by the polymerase efficiency which resulted in 20.83, 16.19, 16.87, and 20.98 total template doubling for the Hi-Fi 2X, Ultra II, KAPA, and SuperFi polymerases, respectively.

### MERIT

SAMtools mpileup was used for variant identification step in the MERIT pipeline. As MERIT was designed for ultra-deep HTS applications, the input options of its SAMtools mpileup were set to accommodate high depths while providing the users the ability to modify these parameters based on each sequencing data’s characteristics. Table S5 in SI Materials summarizes the default input parameters of SAMtools mpileup and those used in MERIT. These parameters ensure the inclusion of all reads and variants for profiling errors, and may not be suitable for calling statistically significant variants.

MERIT extracts the Phred quality score of base substitutions as well as the average Phred quality of bases before and after indels. These quantities are not provided in VCF files generated by SAMtools. Of note, the alternate allele and total depths at indel loci are only accurate in SAMtools’s Pileup files and not in its VCF file. Therefore, to ensure VAF accuracy for both indels and substitutions, the reference and alternate alleles’ depths as well as the total depths for all the variants were extracted from the Pileup file, including all the read in the analysis. Finally, MERIT extracts the position-in-read for all variants. Such information, especially in hybrid-capture sequencing, helps to better quantify the source of errors in HTS platforms.

### Data and code availability

MERIT is open-sourced and available at www.software.khiabanian-lab.org. All the DNA sequencing data used in this study have been deposited in the NCBI’s Sequence Read Archive (SRA) with accession number SRP115798.

## Acknowledgments

The authors gratefully acknowledge the constructive feedback of Alexandra Jacunski. M.H. is a New Jersey Commission on Cancer Research postdoctoral fellow (DFHS17PPC007). H.K. acknowledges support from the American Caner Society (IRG-15-168-01), Rutgers Cancer Institute of New Jersey (P30CA072720), and Rutgers Office of Advanced Research Computing (NIH 1S10OD012346-01A1).

## References

[1] Davide Rossi, Hossein Khiabanian, Valeria Spina, Carmela Ciardullo, Alessio Bruscaggin, Rosella Famà, Silvia Rasi, Sara Monti, Clara Deambrogi, Lorenzo De Paoli, Jiguang Wang, Valter Gattei, Anna Guarini, Robin Foà, Raul Rabadan, and Gianluca Gaidano. Clinical impact of small tp53 mutated subclones in chronic lymphocytic leukemia. Blood, 123(14):2139–2147, 2014.

[2] A. N. Hata, M. J. Niederst, H. L. Archibald, M. Gomez-Caraballo, F. M. Siddiqui, H. E. Mulvey, Y. E. Maruvka, F. Ji, H. E. Bhang, V. Krishnamurthy Radhakrishna, G. Siravegna, H. Hu, S. Raoof, E. Lockerman, A. Kalsy, D. Lee, C. L. Keating, D. A. Ruddy, L. J. Damon, A. S. Crystal, C. Costa, Z. Piotrowska, A. Bardelli, A. J. Iafrate, R. I. Sadreyev, F. Stegmeier, G. Getz, L. V. Sequist, A. C. Faber, and J. A. Engelman. Tumor cells can follow distinct evolutionary paths to become resistant to epidermal growth factor receptor inhibition. Nat Med, 22(3):262–9, 2016.

[3] Edward J Fox, Kate S Reid-Bayliss, Mary J Emond, and Lawrence A Loeb. Accuracy of next generation sequencing platforms. Next generation, sequencing & applications, 1:1000106, 2014.

[4] Isaac Kinde, Jian Wu, Nick Papadopoulos, Kenneth W. Kinzler, and Bert Vogelstein. Detection and quantification of rare mutations with massively parallel sequencing. Proceedings of the National Academy of Sciences, 108(23):9530–9535, 2011.

[5] Daniel Aird, Michael G. Ross, Wei-Sheng Chen, Maxwell Danielsson, Timothy Fennell, Carsten Russ, David B. Jaffe, Chad Nusbaum, and Andreas Gnirke. Analyzing and minimizing pcr amplification bias in illumina sequencing libraries. Genome Biology, 12(2):R18, 2011.

[6] Christopher Huptas, Siegfried Scherer, and Mareike Wenning. Optimized illumina pcr-free library preparation for bacterial whole genome sequencing and analysis of factors influencing de novo assembly. BMC Research Notes, 9:269, 2016.

[7] Jeffrey A. Martin and Zhong Wang. Next-generation transcriptome assembly. Nat Rev Genet, 12(10):671–682, 10 2011.

[8] Michael W Schmitt, Scott R Kennedy, Jesse J Salk, Edward J Fox, Joseph B Hiatt, and Lawrence A Loeb. Detection of ultra-rare mutations by next-generation sequencing. Proceedings of the National Academy of Sciences of the United States of America, 109(36):14508–14513, 09 2012.

[9] Justin Jee, Aviram Rasouly, Ilya Shamovsky, Yonatan Akivis, Susan R. Steinman, Bud Mishra, and Evgeny Nudler. Rates and mechanisms of bacterial mutagenesis from maximum-depth sequencing. Nature, 534(7609):693–696, 06 2016.

[10] Tanja Magoč and Steven L Salzberg. Flash: fast length adjustment of short reads to improve genome assemblies. Bioinformatics, 27(21):2957–2963, 11 2011.

[11] Binghang Liu, Jianying Yuan, Siu-Ming Yiu, Zhenyu Li, Yinlong Xie, Yanxiang Chen, Yujian Shi, Hao Zhang, Yingrui Li, Tak-Wah Lam, and Ruibang Luo. Cope: an accurate k-mer-based pair-end reads connection tool to facilitate genome assembly. Bioinformatics, 28(22):2870, 2012.

[12] Sébastien Rodrigue, Arne C Materna, Sonia C Timberlake, Matthew C Blackburn, Rex R Malmstrom, Eric J Alm, and Sallie W Chisholm. Unlocking short read sequencing for metagenomics. PLoS ONE, 5(7):e11840, 2010.

[13] Kensuke Nakamura, Taku Oshima, Takuya Morimoto, Shun Ikeda, Hirofumi Yoshikawa, Yuh Shiwa, Shu Ishikawa, Margaret C. Linak, Aki Hirai, Hiroki Takahashi, Md. Altaf-Ul-Amin, Naotake Ogasawara, and Shigehiko Kanaya. Sequence-specific error profile of illumina sequencers. Nucleic Acids Research, 39(13):e90, 2011.

[14] Wei Shao, Valerie F. Boltz, Jonathan E. Spindler, Mary F. Kearney, Frank Maldarelli, John W. Mellors, Claudia Stewart, Natalia Volfovsky, Alexander Levitsky, Robert M. Stephens, and John M. Coffin. Analysis of 454 sequencing error rate, error sources, and artifact recombination for detection of low-frequency drug resistance mutations in hiv-1 dna. Retrovirology, 10(1):18, 2013.

[15] Johanna Brodin, Mattias Mild, Charlotte Hedskog, Ellen Sherwood, Thomas Leitner, Björn Andersson, and Jan Albert. Pcr-induced transitions are the major source of error in cleaned ultra-deep pyrosequencing data. PLoS ONE, 8(7):e70388, 2013.

[16] Juliane C Dohm, Claudio Lottaz, Tatiana Borodina, and Heinz Himmelbauer. Substantial biases in ultra-short read data sets from high-throughput dna sequencing. Nucleic Acids Research, 36(16):e105–e105, 09 2008.

[17] Melanie Schirmer, Umer Z. Ijaz, Rosalinda D’Amore, Neil Hall, William T. Sloan, and Christopher Quince. Insight into biases and sequencing errors for amplicon sequencing with the illumina miseq platform. Nucleic Acids Research, 2015.

[18] Aaron McKenna, Matthew Hanna, Eric Banks, Andrey Sivachenko, Kristian Cibulskis, Andrew Kernytsky, Kiran Garimella, David Altshuler, Stacey Gabriel, Mark Daly, and Mark A DePristo. The genome analysis toolkit: A mapreduce framework for analyzing next-generation dna sequencing data. Genome Research, 20(9):1297–1303, 09 2010.

[19] Heng Li, Bob Handsaker, Alec Wysoker, Tim Fennell, Jue Ruan, Nils Homer, Gabor Marth, Goncalo Abecasis, Richard Durbin, and 1000 Genome Project Data Processing Subgroup. The sequence alignment/map format and samtools. Bioinformatics, 25(16):2078–2079, 08 2009.

[20] Erik Garrison and Gabor Marth. Haplotype-based variant detection from short-read sequencing. arXiv:1207.3907 [q-bio.GN], 2012.

[21] Xiangtao Liu, Shizhong Han, Zuoheng Wang, Joel Gelernter, and Bao-Zhu Yang. Variant callers for next-generation sequencing data: A comparison study. PLOS ONE, 8(9), 09 2013.

[22] Sohyun Hwang, Eiru Kim, Insuk Lee, and Edward M. Marcotte. Systematic comparison of variant calling pipelines using gold standard personal exome variants. Scientific Reports, 5:17875 EP —, 12 2015.

[23] Anne Bruun Krøigård, Mads Thomassen, Anne-Vibeke Lænkholm, Torben A Kruse, and Martin Jakob Larsen. Evaluation of nine somatic variant callers for detection of somatic mutations in exome and targeted deep sequencing data. PLoS ONE, 11(3):e0151664, 2016.

[24] Andre P Masella, Andrea K Bartram, Jakub M Truszkowski, Daniel G Brown, and Josh D Neufeld. Pandaseq: paired-end assembler for illumina sequences. BMC Bioinformatics, 13:31–31, 2012.

[25] Jiajie Zhang, Kassian Kobert, Tomáš Flouri, and Alexandros Stamatakis. Pear: a fast and accurate illumina paired-end read merger. Bioinformatics, 30(5):614–620, 03 2014.

[26] Mizuki Ohno, Kunihiko Sakumi, Ryutaro Fukumura, Masato Furuichi, Yuki Iwasaki, Masaaki Hokama, Toshimichi Ikemura, Teruhisa Tsuzuki, Yoichi Gondo, and Yusaku Nakabeppu. 8-oxoguanine causes spontaneous de novo germline mutations in mice. Scientific Reports, 4:4689 EP -, 04 2014.

[27] K C Cheng, D S Cahill, H Kasai, S Nishimura, and L A Loeb. 8-hydroxyguanine, an abundant form of oxidative dna damage, causes g 1 and a c substitutions. Journal of Biological Chemistry, 267(1):166–172, 1992.

[28] Maura Costello, Trevor J Pugh, Timothy J Fennell, Chip Stewart, Lee Lichtenstein, James C Meldrim, Jennifer L Fostel, Dennis C Friedrich, Danielle Perrin, Danielle Dionne, Sharon Kim, Stacey B Gabriel, Eric S Lander, Sheila Fisher, and Gad Getz. Discovery and characterization of artifactual mutations in deep coverage targeted capture sequencing data due to oxidative dna damage during sample preparation. Nucleic Acids Research, 41(6):e67–e67, 04 2013.

[29] Patricia A. J. Muller and Karen H. Vousden. p53 mutations in cancer. Nature cell biology, 15(1):2–8, 01 2013. Copyright - Copyright Nature Publishing Group Jan 2013; Last updated - 2014–06-15.

[30] Rachel B. Darman, Michael Seiler, Anant A. Agrawal, Kian H. Lim, Shouyong Peng, Daniel Aird, Suzanna L. Bailey, Erica B. Bhavsar, Betty Chan, Simona Colla, Laura Corson, Jacob Feala, Peter Fekkes, Kana Ichikawa, Gregg F. Keaney, Linda Lee, Pavan Kumar, Kaiko Kunii, Crystal MacKenzie, Mark Matijevic, Yoshiharu Mizui, Khin Myint, Eun Sun Park, Xiaoling Puyang, Anand Selvaraj, Michael P. Thomas, Jennifer Tsai, John Y. Wang, Markus Warmuth, Hui Yang, Ping Zhu, Guillermo Garcia-Manero, Richard R. Furman, Lihua Yu, Peter G. Smith, and Silvia Buonamici. Cancer-associated {SF3B1} hotspot mutations induce cryptic 3’ splice site selection through use of a different branch point. Cell Reports, 13(5):1033–1045, 2015.

[31] Lili Wang, Angela N. Brooks, Jean Fan, Youzhong Wan, Rutendo Gambe, Shuqiang Li, Sarah Hergert, Shanye Yin, Samuel S. Freeman, Joshua Z. Levin, Lin Fan, Michael Seiler, Silvia Buonamici, Peter G. Smith, Kevin F. Chau, Carrie L. Cibulskis, Wandi Zhang, Laura Z. Rassenti, Emanuela M. Ghia, Thomas J. Kipps, Stacey Fernandes, Donald B. Bloch, Dylan Kotliar, Dan A. Landau, Sachet A. Shukla, Jon C. Aster, Robin Reed, David S. DeLuca, Jennifer R. Brown, Donna Neuberg, Gad Getz, Kenneth J. Livak, Matthew M. Meyerson, Peter V. Kharchenko, and Catherine J. Wu. Transcriptomic characterization of {SF3B1} mutation reveals its pleiotropic effects in chronic lymphocytic leukemia. Cancer Cell, 30(5):750–763, 2016.

[32] David Sims, Ian Sudbery, Nicholas E. Ilott, Andreas Heger, and Chris P. Ponting. Sequencing depth and coverage: key considerations in genomic analyses. Nat Rev Genet, 15(2):121–132, 02 2014.

[33] New England BioLabs, Inc. Nebnext high-fidelity 2x pcr master mix. http://www.international.neb.com, 2017. Accessed: 2017–07-06.

[34] New England BioLabs, Inc. Nebnext ultra ii q5 master mix. http://www.international.neb.com, 2017. Accessed: 2017–07-06.

[35] Kapa Biosystems. Kapa hifi pcr kits. http://www.kapabiosystems.com, 2017. Accessed: 2017–07-06.

[36] Thermo Fisher Scientific. Invitrogen platinum superfi dna polymerase. http://www.thermofisher.com, 2017. Accessed: 2017–07-06.

[37] K A Eckert and T A Kunkel. Dna polymerase fidelity and the polymerase chain reaction. Genome Research, 1(1):17–24, 1991.

[38] N F Cariello, J A Swenberg, and T R Skopek. Fidelity of thermococcus litoralis dna polymerase (vent) in pcr determined by denaturing gradient gel electrophoresis. Nucleic Acids Research, 19(15):4193–4198, 08 1991.

[39] Peter McInerney, Paul Adams, and Masood Z. Hadi. Error rate comparison during polymerase chain reaction by dna polymerase. Molecular Biology International, 12014:1–8, 2014.

[40] Dmitriy A. Shagin, Irina A. Shagina, Andrew R. Zaretsky, Ekaterina V. Barsova, Ilya V. Kelmanson, Sergey Lukyanov, Dmitriy M. Chudakov, and Mikhail Shugay. A high-throughput assay for quantitative measurement of pcr errors. Scientific Reports, 7(1):2718, 2017.

[41] Vladimir Potapov and Jennifer L. Ong. Examining sources of error in pcr by single-molecule sequencing. PLOS ONE, 12(1):1–19, 01 2017.

[42] Matthew S. Hestand, Jeroen Van Houdt, Francesca Cristofoli, and Joris R. Vermeesch. Polymerase specific error rates and profiles identified by single molecule sequencing. Mutation Research/Fundamental and Molecular Mechanisms of Mutagenesis, 784–785:39 — 45, 2016.

[43] Iwanka Kozarewa, Zemin Ning, Michael A Quail, Mandy J Sanders, Matthew Berriman, and Daniel J Turner. Amplification-free illumina sequencing-library preparation facilitates improved mapping and assembly of (g+c)-biased genomes. Nat Meth, 6(4):291–295, 04 2009.

[44] Lira Mamanova, Robert M Andrews, Keith D James, Elizabeth M Sheridan, Peter D Ellis, Cordelia F Langford, Tobias W B Ost, John E Collins, and Daniel J Turner. Frt-seq: amplification-free, strand-specific transcriptome sequencing. Nat Meth, 7(2):130–132, 02 2010.

[45] Dianne I Lou, Jeffrey A Hussmann, Ross M McBee, Ashley Acevedo, Raul Andino, William H Press, and Sara L Sawyer. High-throughput dna sequencing errors are reduced by orders of magnitude using circle sequencing. Proceedings of the National Academy of Sciences of the United States of America, 110(49):19872–19877, 12 2013.

[46] M. T. Gregory, J. A. Bertout, N. G. Ericson, S. D. Taylor, R. Mukherjee, H. S. Robins, C. W. Drescher, and J. H. Bielas. Targeted single molecule mutation detection with massively parallel sequencing. Nucleic Acids Res, 44(3):e22, 2016.

[47] Mario Cazzola, Marianna Rossi, Luca Malcovati, and on behalf of the Associazione Italiana per la Ricerca sul Cancro Gruppo Italiano Malattie Mieloproliferative. Biologic and clinical significance of somatic mutations of sf3b1 in myeloid and lymphoid neoplasms. Blood, 121(2):260–269, 01 2013.

[48] D. Rossi, H. Khiabanian, V. Spina, C. Ciardullo, A. Bruscaggin, R. Fama, S. Rasi, S. Monti, C. Deambrogi, L. De Paoli, J. Wang, V. Gattei, A. Guarini, R. Foa, R. Rabadan, and G. Gaidano. Clinical impact of small tp53 mutated subclones in chronic lymphocytic leukemia. Blood, 123(14):2139–47, 2014.

[49] S. Rasi, H. Khiabanian, C. Ciardullo, L. Terzi-di Bergamo, S. Monti, V. Spina, A. Bruscaggin, M. Cerri, C. Deambrogi, L. Martuscelli, A. Biasi, E. Spaccarotella, L. De Paoli, V. Gattei, R. Foa, R. Rabadan, G. Gaidano, and D. Rossi. Clinical impact of small subclones harboring notch1, sf3b1 or birc3 mutations in chronic lymphocytic leukemia. Haematologica, 101(4):e135–8, 2016.

[50] Ferran Nadeu, Julio Delgado, Cristina Royo, Tycho Baumann, Tatjana Stankovic, Magda Pinyol, Pedro Jares, Alba Navarro, David Martín-García, Sílvia Beà, Itziar Salaverria, Ceri Oldreive, Marta Aymerich, Helena Suárez-Cisneros, Maria Rozman, Neus Villamor, Dolors Colomer, Armando López-Guillermo, Marcos González, Miguel Alcoceba, Maria José Terol, Enrique Colado, Xose S Puente, Carlos López-Otín, Anna Enjuanes, and Elϭas Campo. Clinical impact of clonal and subclonal tp53, sf3b1, birc3, notch1, and atm mutations in chronic lymphocytic leukemia. Blood, 127(17):2122–2130, 04 2016.

[51] Raul Rabadan, Sonia Marsilio, Nicholas Chiorazzi, Laura Pasqualucci, and Hossein Khiabanian. Highly sensitive detection of small variants in multi-sample ultra-deep tumor sequencing. bioRxiv, 2017.

[52] Andreas Untergasser, Ioana Cutcutache, Triinu Koressaar, Jian Ye, Brant C Faircloth, Maido Remm, and Steven G Rozen. Primer3—new capabilities and interfaces. Nucleic Acids Research, 40(15):e115–e115, 08 2012.

[53] Heng Li and Richard Durbin. Fast and accurate short read alignment with burrows-wheeler transform. Bioinformatics, 25(14):1754–1760, 07 2009.

